# Sequencing ultra-rare targets with compound nucleic acid cytometry

**DOI:** 10.1101/2020.09.01.278275

**Authors:** Chen Sun, Kai-Chun Chang, Adam R. Abate

## Abstract

Targeted sequencing enables sensitive and cost-effective analysis by focusing resources on molecules of interest. Existing methods, however, are limited in enrichment power and target capture length. Here, we present a novel method that uses compound nucleic acid cytometry to achieve million-fold enrichments of molecules >10 kbp in length using minimal prior target information. We demonstrate the approach by sequencing HIV proviruses in infected individuals. Our method is useful for rare target sequencing in research and clinical applications, including for identifying cancer-associated mutations or sequencing viruses infecting cells.

## Introduction

Target enrichment focuses valuable sequencing on important molecules and is useful when the sample comprises a large background of uninteresting DNA^1^. For instance, characterizing HIV genomic diversity is important for understanding persistent infection, but under treatment viral DNA is outnumbered by human DNA by billions of times^2-4^. In metagenomic analyses, organisms of interest may be present at a few percent^5-7^, while in human genetic disease, variants may be present at fractions of a percent^8-10^. In instances like these, sequencing all DNA is wasteful because only a fraction of reads corresponds to the region of interest. The most common target enrichment strategies are based on PCR amplification or hybridization capture^1,11,12^. PCR methods recover only the amplified portion and miss information beyond primers^13,14^. Hybridization capture recovers information extending beyond probes, but can require hundreds of probes^15,16^; this necessitates considerable prior information for probe design, which is often unavailable, especially when little information is known about the region of interest, such as in novel microbe or genetic lesion sequencing^17,18^.

Nucleic acid cytometry (NAC) is a conceptually novel approach to target enrichment based on droplet microfluidics^19^. The overarching principle is to physically isolate molecules by hydrodynamic sorting. Target identification is accomplished using droplet PCR, while isolation is accomplished by sorting positive droplets^19- 22^. The approach is akin to querying a diverse mixture for keyword subsequences, and isolating all molecules containing the keyword. The critical factor in NAC enrichment is sensitivity for recovering the target of interest. Sensitivity, in turn, is limited by the number of droplet PCRs that can be sorted which, presently, is ∼10 million. Considering losses in DNA recovery and the need for sufficient material to perform sequencing, current enrichments are capped to ∼30,000, allowing NAC to maximally concentrate the target by this factor^5,8,23,24^. This enrichment is insufficient for applications with ultra-rare targets below one in a million. To broaden the applicability of NAC, a strategy to increase enrichment power is needed.

In this study, we demonstrate the ability to perform NAC repeatedly on a sample to achieve compound enrichment over multiple rounds. The final enrichment is the product of each round, allowing a ∼6 million-fold enrichment over two rounds. This is ∼200-fold higher than enrichments with the next best technology^8,15,23^. To demonstrate the approach, we use it to isolate and sequence single HIV genomes from infected individuals. No other enrichment approach has the sensitivity to recover and sequence such rare single virus genomes. Compound NAC provides a general platform for recovering long, ultra-rare molecules with minimal prior sequence information.

### Overview

Nucleic acid cytometry isolates molecules of interest from a mixed population based on specific sequence biomarkers^19^. This is achieved by combining droplet TaqMan PCR identification and microfluidic droplet sorting to physically isolate molecules based on the TaqMan signal (Fig. 1a). The DNA mixture is partitioned at limiting dilution such that individual droplets rarely contain more than one target. The purity of the target sequence is *N*_*T*_/ *N*_*T*_ + *N*_*O*_ before sorting, and *N*_*T*_*D*/ (*N*_*T*_ + *N*_*O*_) *N*_*T*_ + *f D* after, where *N*_*T*_ is the number of target molecules (positively sorted drops), *N*_*O*_ the total number of off-target molecules, *D* the total number of droplets and *f* the assay false positive rate. Thus, the enrichment power is (*N*_*T*_/*D* + *f*)^+,^ (derivation in supplementary material). Because the false positive rate is generally small (∼10^−5^) and difficult to reduce^25^, the best way to increase enrichment power is to encapsulate the sample into more droplets, which thus delivers fewer co-encapsulated off-target molecules per sorted positive. However, the number of droplets that can be sorted is limited to ∼10 million^8,19^. Consequently, the maximum practical enrichment that can be achieved per NAC round is ∼10^4^.

**Figure 1.**
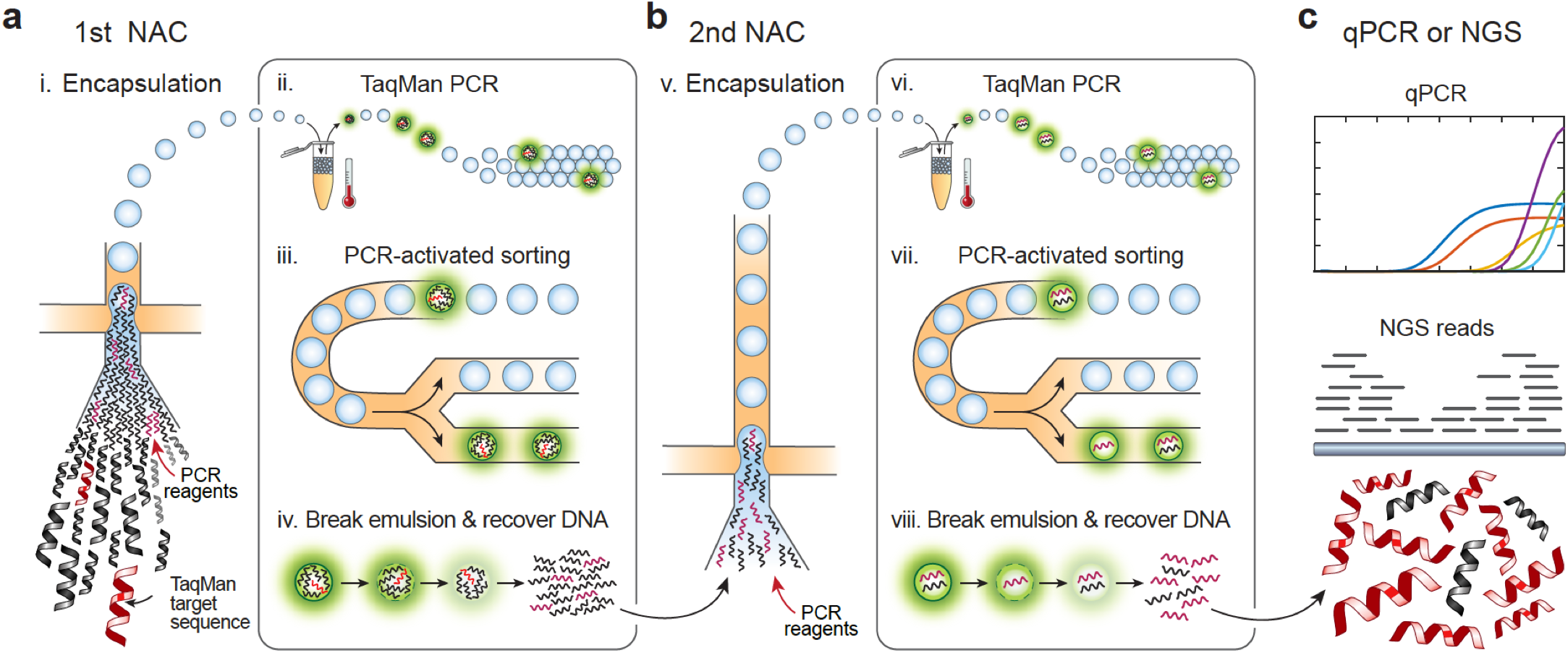
Schematic of compound NAC workflow. **(a)** A mixed DNA sample is sorted in a first NAC round using TaqMan targeting a desired sequence biomarker. Each NAC round comprises (i) DNA encapsulation with TaqMan reagents; (ii) in-droplet PCR to generate fluorescence when the target is present; (iii) sorting to select positive drops; (iv) recovery of sorted DNA by droplet demulsification. **(b)** DNA recovered from the first NAC round is diluted and processed through another round consisting of the same steps (v-viii). **(c)** The double-enriched DNA is analyzed by qPCR to estimate enrichment, and sequenced.

Like hybridization capture, NAC does not fragment the original target molecules. However, in contrast to hybridization capture, NAC can recover long intact targets (>100 kbp) present in a sample over a wide range of DNA concentrations^8^. These features allow it to be performed repeatedly on a sample such that the enrichments compound (Fig. 1b). In such a strategy, the overall enrichment with two rounds is [(*N*_*T*_/*D*_1_ + *f*_1_)(*N*_*T*_/*D*_2_ + *f*_2_)]^+,^ when using *D*_1_ drops in the first round and *D*_2_ drops in the second. Compound enrichment thus allows marked increases to enrichment compared to sorting more drops in a single round. For example, for a total of ∼10^7^ drops sorted, one round typically achieves ∼10^3^ enrichment of 10,000 target molecules, while two consecutive rounds achieve ∼10^6^. Obtaining such an enrichment with a single-round of NAC would require sorting over a billion droplets, which is impractical. The resultant concentrated DNA is intact and readily amenable to qPCR or sequencing analysis (Fig. 1c).

### Microfluidic workflow for compound enrichment

NAC uses ultrahigh-throughput microfluidics to perform, analyze, and sort millions of PCR reactions. Flow focusing loads sample DNA with TaqMan reagents in ∼45 μm droplets at ∼2.5 kHz, partitioning the entire 150 μL reaction in ∼3 million drops in ∼20 min (Fig. 2a). The drops are thermocycled, generating TaqMan fluorescence when the target is present (Fig. 2b). The drops are analyzed and sorted using a laser-induced fluorescence detector and dielectrophoretic droplet deflector^26,27^ (Fig. 2c). We operate this integrated device at ∼400 Hz to ensure accurate sorting and efficient positive recovery, screening ∼3 million drops in ∼2 hr. The sorted target molecules are recovered by droplet demulsification with perfluoro-octanol^27^ and diluted into new TaqMan reagents for the next round of NAC.

**Figure 2.**
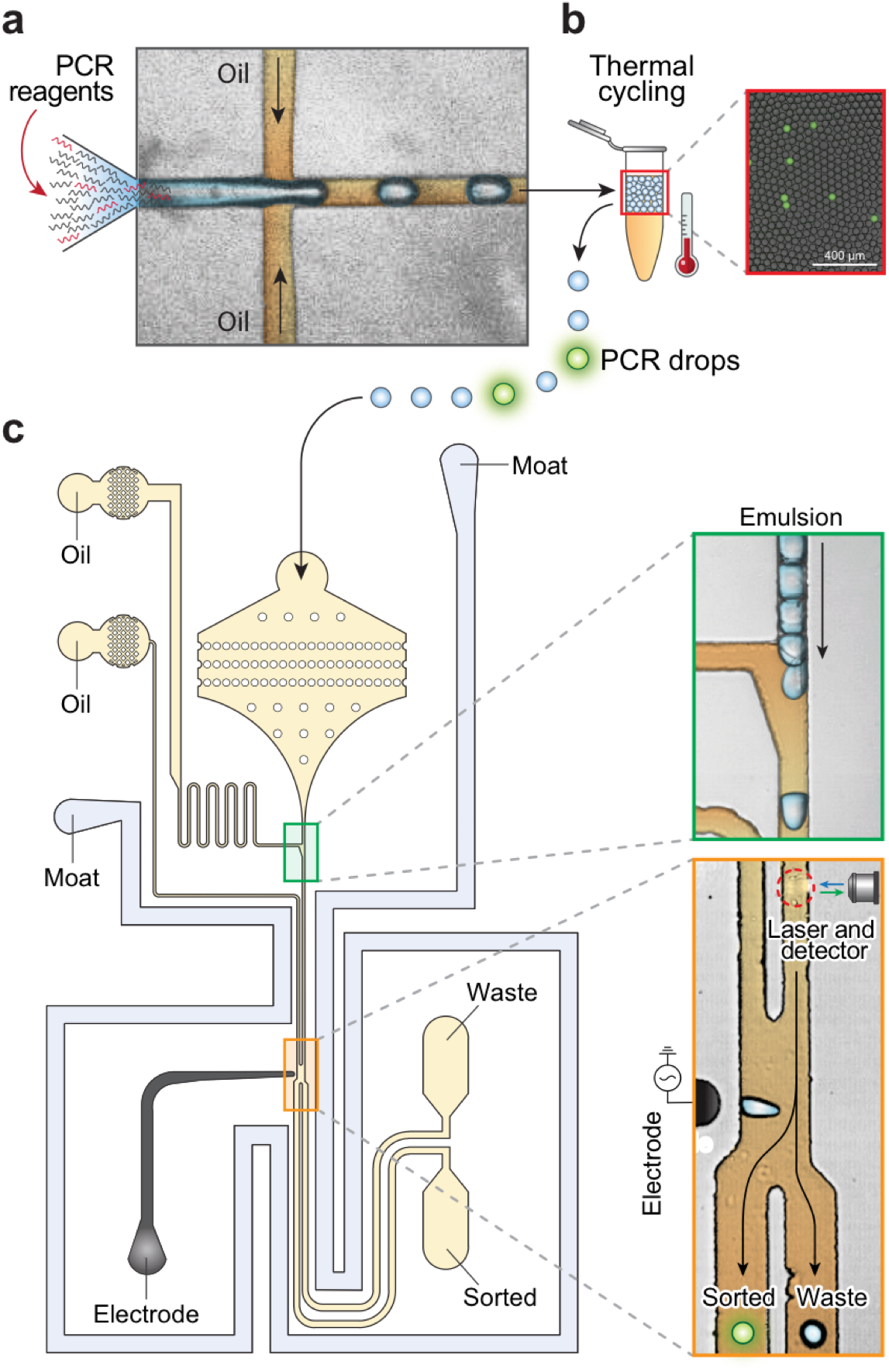
Microfluidic devices of NAC. **(a)** Droplet encapsulation of DNA and PCR reagents. **(b)** In-drop PCR to generate fluorescence when the target is present. The merged bright field/fluorescence image shows a representative sample post thermocycling. **(c)** Fluorescence-activated sorting selects droplets containing target sequences. Scale bars: 400 µm.

### Compound enrichment of ΦX 174 virus

To demonstrate the power of compound NAC, we apply it to enrich ΦX 174 viral genomes from a 10^7^-fold greater background of lambda DNA. The TaqMan set used in each round detects a different region of the ΦX 174 genome, preventing amplicons carried over from the first round generating false positives in the second (Fig. 3a). Both sets reliably detect ΦX 174 DNA (Supplementary Fig. 1). To obtain an optimal enrichment of ∼400, we set the target concentration such that in the first round 0.24% of droplets are positive (Fig. 3b(i)). The recovered DNA is diluted into fresh reaction buffer again to achieve another 400-fold enrichment, and subjected to another round of NAC (Fig. 3b(ii)). Because the method is nondestructive, the number of positive drops should be equal for both rounds, but sample loss during preparation for the second results in slightly fewer total positives (Fig. 3b(iii)). To confirm enrichment, we use qPCR to measure the fractions of ΦX 174 and lambda DNA in the sorted samples. After a single round, the qPCR curve for ΦX 174 shifts to lower cycles (concentrated) while that for lambda shifts to higher cycles (diluted), illustrating enrichment (Fig. 3c(i)). For two rounds compounded, these shifts are greater (Fig. 3c(ii)). To quantify the enrichments, we calculate the enrichment factor *e* based on the cross-threshold values of the qPCR curves^20^. For one round of sorting ∼3 million droplets, the estimated enrichment is ∼150. For two rounds of sorting comprising a total of ∼6 million droplets, the enrichment is ∼16,000. To achieve this enrichment in one round would require sorting ∼300 million droplets, totaling 16 mL of PCR reagent, and a week of nonstop sorting.

**Figure 3.**
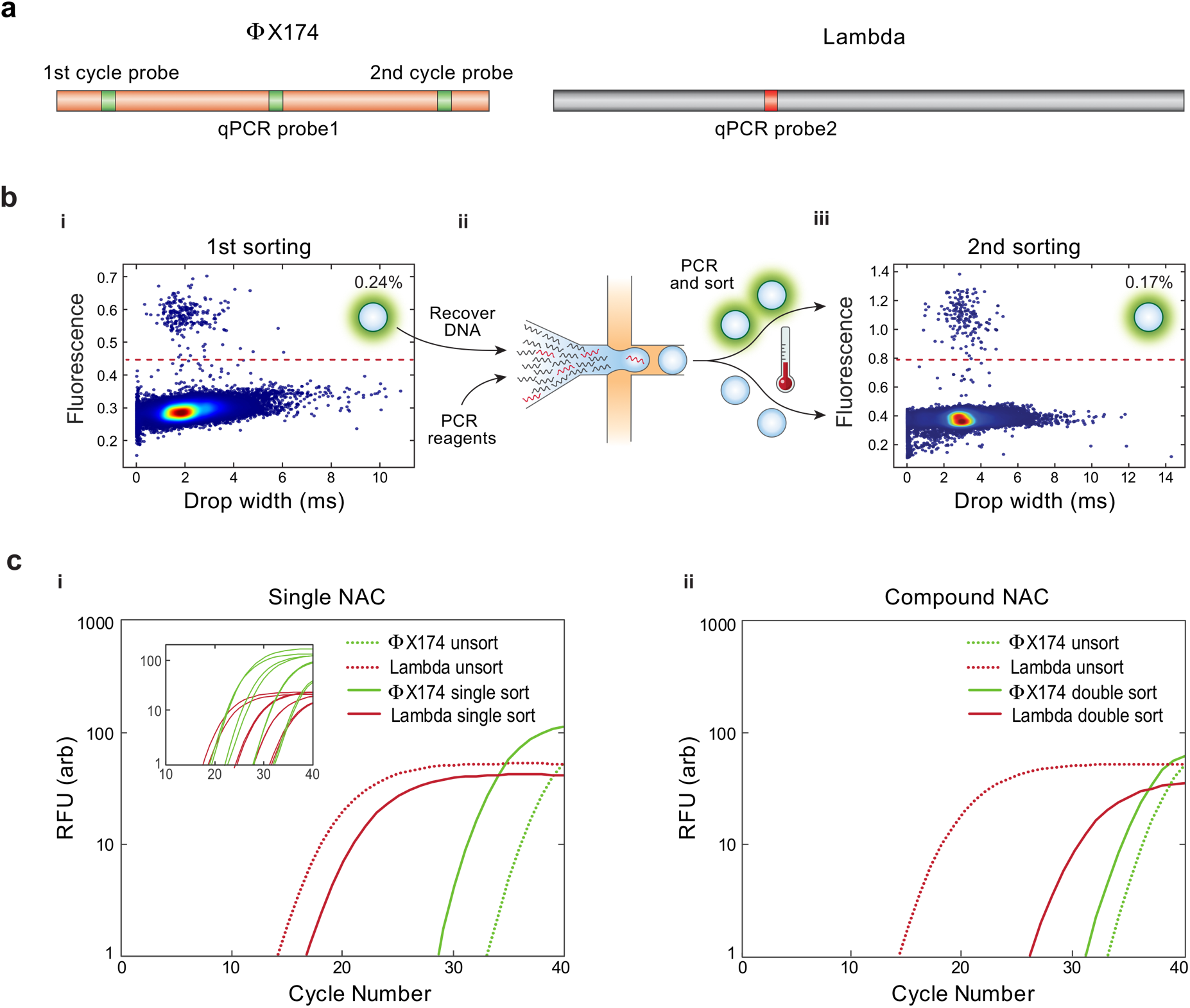
Enrichment of ΦX 174 DNA from a background of Lambda DNA with compound NAC. **(a)** TaqMan assays detect droplets containing ΦX 174 (green) and Lambda (red) DNA. **(b)** The microfluidic sorter interrogates the droplets for fluorescence and sorts PCR positives. (i) Scatter plot of fluorescence versus size of drops from first NAC round, with 0.24% positive. (ii) DNA from the first round is recovered, diluted, and processed again. (iii) Scatter plot of fluorescence versus size of drops from the second NAC round, with 0.17% positive. **(c)** qPCR plots for (i) single and (ii) double-enriched DNA; based on curve shifts, single-round sorting enriches ΦX 174 by ∼150-fold, and double-round sorting by ∼16,000-fold. Inset in (i) shows ΦX 174 and Lambda standard curves.

### Single genome sequencing of ultra-rare HIV proviruses

During effective antiretroviral therapy, HIV persists in a latent state and circulates at extremely low levels, with human DNA outnumbering it by over a billion-fold^2,3,28^. Under such circumstances, unbiased sequencing would recover a minute fraction of one viral genome per human genome sequenced. To obtain comprehensive information on the genetics of HIV under such circumstances thus requires potent enrichment of the virus. The only effective strategy presently available is terminal dilution PCR in well plates^3,4,29^. This brute force approach aliquots thousands of cells in hundreds of microwells, using long ranged, multi-primer amplification to obtain near full-length HIV genomes. However, in addition to often generating artifacts that can confound analysis, the approach does not obtain the crucial virus-host junction with the complete virus genome in a single contig. Without this dual information, specific proviruses cannot be related to host insertion sites and thereby the combination associated to disease behavior^3,30^. Consequently, other strategies must be employed to infer viral genome and host junction relationships^29,30^.

Due to the potent enrichment enabled by compound NAC and its ability to recover intact DNA fragments spanning integrated HIV genomes, such analyses are possible. To demonstrate this, we use compound NAC to isolate and sequence HIV proviruses from a patient infected cell expansion^28^. This clonal lineage of infected cells contains one HIV provirus per ∼100 cells, each bearing the same provirus. To demonstrate the enrichment power of compound NAC, we dilute the sample with non-HIV cell gDNA at a 1:30 ratio. This dilution models the concentration of latent infection. Due to the extreme rarity of HIV DNA in this sample, we use a multiplexed TaqMan PCR targeting multiple conserved regions of HIV in the second cycle for accurate detection and isolation before sequencing (Fig. 4a). The DNA mixture is encapsulated in droplets and ones containing HIV genomes are isolated. By incorporating in-droplet whole genome amplification before the second round^31^, each sorted droplet yields ∼3 pg DNA, just enough for sequencing (Fig. 4b). This novel workflow affords superior enrichment and single drop sequencing, allowing recovery of integrated provirus genomes (Fig.4c, first to third row). We thus identify the integration site in host gene ARIH2 by extracting virus-human chimeric reads from the sequencing data. Shearing during PCR and DNA preparation results in partial genome dropout. Thus, by assembling reads from 3 droplets, we obtain complete coverage of the full-length viral genome (Fig.4c, 4th row). In total, sequencing after two rounds detects ∼6 million times more proviral reads compared to the initial sample. These results illustrate that compound NAC enables sequencing of extremely rare HIV proviruses, and that the enriched molecules retain information on the genetic context of the integration.

**Figure 4.**
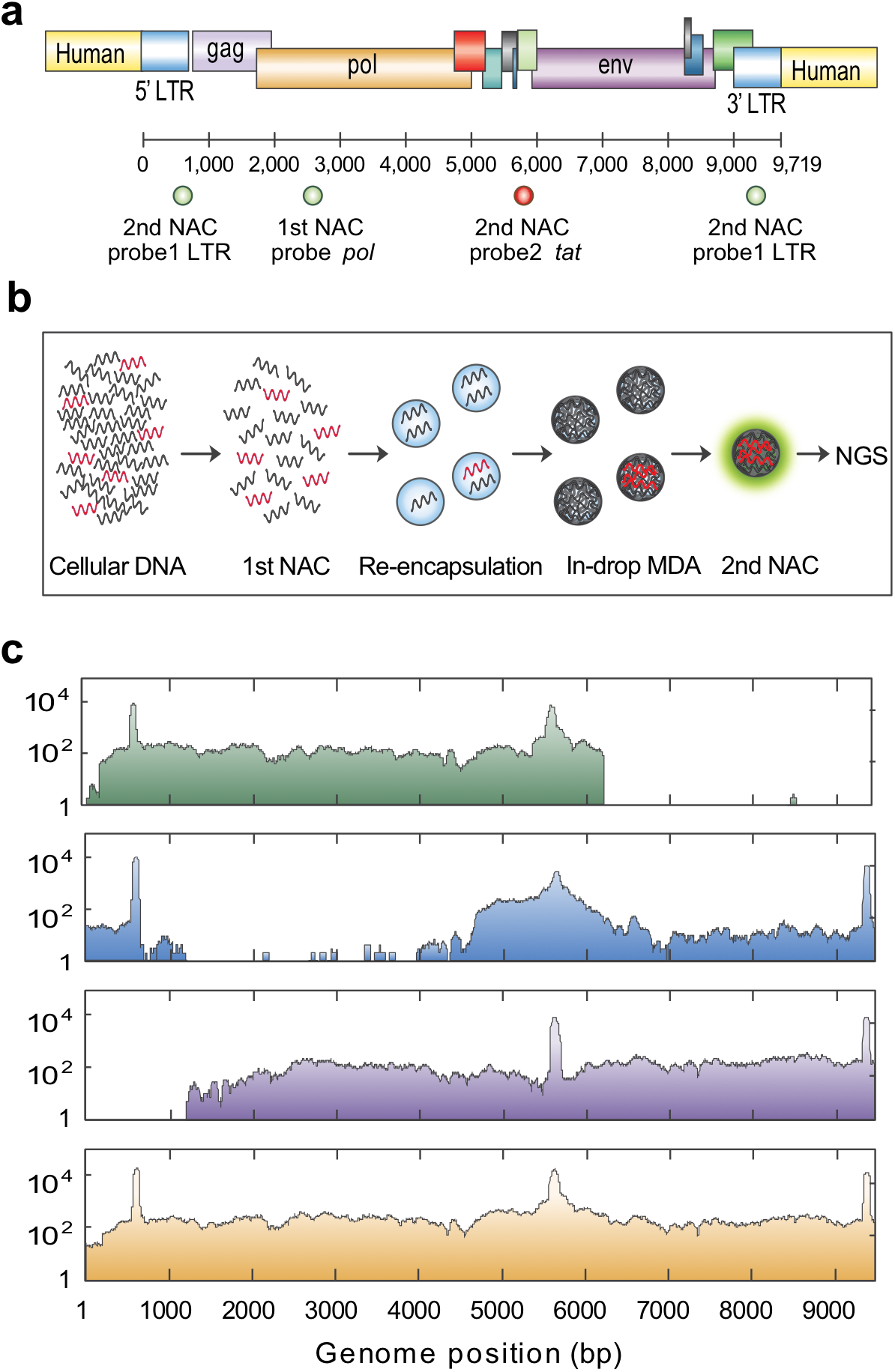
Compound enrichment and sequencing of HIV proviruses in infected individuals. **(a)** The first round of NAC uses a TaqMan assay targeting the *pol* gene (FAM). The second round of NAC uses a degenerate TaqMan assay targeting the long terminal repeat (LTR, FAM) and *tat* gene (Cy5). **(b)** Implementation of in-droplet multiple-displacement amplification before the second round generates sufficient material for single droplet sequencing. **(c)** HIV genome and integration site coverage maps for three sorted drops. Aggregating all data assembles the full-length HIV genome including the human integration site. The peaks at LTR and *tat* are the TaqMan PCR amplicons.

## Discussion

By leveraging droplet microfluidics, NAC enables enrichment of target molecules containing sequence biomarkers. By processing the sample repeatedly, feeding the output of the first round into the input of the second, enrichment compounds, allowing recovery of ultra-rare targets. In single-round NAC, maximum enrichment is limited by the false positive droplet rate. Partitioning the sample into more droplets enhances enrichment in a linear fashion, but does not allow the marked increases required for ultra-rare targets. Moreover, such brute force also increases cost and processing time and becomes impractical beyond enrichments of 30,000^8,23^. Our method eliminates the need for large numbers of droplets and increases maximum enrichment to 10^9^ fold, allowing highly specific target recovery.

Compound NAC allows isolation of long molecules with minimal prior sequence information, opening new avenues in target enrichment. For example, million-fold enrichment of >100 kbp molecules is useful for a variety of ultra-rare target applications, including characterizing novel human genetic mutations or natural product gene clusters in metagenomic samples^5,8^. Additionally, the approach is generalizable to other targets because it uses TaqMan PCR to define the sequence biomarker of capture and, thus, can be applied to any nucleic acid detectable by this assay, including RNA by adding a reverse transcription step; this would allow sequencing of fusion genes or low-abundance variants. Finally, as we have shown, implementation of in-droplet MDA allows sequencing of compound enriched single molecules, making it a powerful tool for single virus genomics.

## Methods

### Microfluidic device fabrication

The microfluidic devices were fabricated in Polydimethylsiloxane (PDMS) using standard soft lithography. Photomasks designed by AutoCAD were printed on transparencies and the features on the photomask transferred to a silicon wafer (University Wafer) using negative photoresist (MicroChem, SU-8 2025) by UV photolithography. PDMS (Dow Corning, Sylgard 184) prepolymer mixture of polymer and cross-linker at a ratio of 10:1 was poured over the pattered silicon wafer and cured in a 65 °C oven for 2 hr. PDMS replica was peeled off and punched for inlets and outlets by a 0.75 mm biopsy core (World Precision Instruments). The PDMS slab was bound to a clean glass using an oxygen plasma cleaner (Harrick Plasma), followed by baking at 65 °C for 30 min to ensure strong bonding between the PDMS and glass. The microfluidic channels were treated with Aquapel (PPG Industries) and baked at 65 °C overnight for hydrophobicity.

### Droplet TaqMan PCR

ΦX 174 virion DNA and Lambda DNA (New England BioLabs) were added to PCR reagents containing 1X Platinum Multiplex PCR Master Mix (Life Technologies, catalog no. 4464269), 200 nM TaqMan probe (IDT), 1 μM forward primer and 1 μM reverse primer (IDT), 2.5% (w/w) Tween® 20 (Fisher Scientific), 2.5% (w/w) Poly(ethylene glycol) 6000 (Sigma-Aldrich) and 0.8 M 1,2-propanediol (Sigma-Aldrich). Tween® 20 and Poly(ethylene glycol) 6000 were used to increase stability of droplets during thermal cycling^22^. 1,2-propanediol was used as a PCR enhancer when low temperature was used for denaturation^32^. Two syringes backfilled with HFE-7500 fluorinated oil (3M, catalog no. 98-0212-2928-5) were loaded with (1) TaqMan PCR reaction mix, (2) HFE-7500 oil with 2% (w/w) PEG-PFPE amphiphilic block copolymer surfactant (RNA biotechnologies, catalog no. 008-FluoroSurfactant-1G). The aqueous phase and oil phase were injected into a flow-focus droplet maker at controlled flow rates (400 μl/h for PCR mix and 800 μl/h for oil phase) sustained by computer programmed syringe pumps (New Era). Monodispersed droplets (diameter ∼40 μm) were generated and collected to PCR tubes via polyethylene tubing. The bottom oil was then removed and replaced with FC-40 fluorinated oil (Sigma-Aldrich, catalog no. 51142-49-5) with 5% (w/w) PEG-PFPE amphiphilic block copolymer surfactant for better droplet stability before putting the emulsion into a thermal cycler (Bio-Rad, T100 model). Thermal cycling was performed at: 2 min 30 s at 86 °C; 35 cycles of 30 s at 86 °C, 1 min 30 s at 60 °C and 30 s at 72 °C; and a final extension of 5 min at 72 °C. A low denaturation temperature of 86 °C was used to minimize DNA fragmentation. After PCR, a small aliquot of drops was visualized with an EVOS inverted fluorescence microscope. Another small aliquot of drops was taken and broken with 10% (v/v) solution of perfluoro-octanol (Sigma-Aldrich, catalog no. 370533) and addition of 10 μl DI water, followed by gentle vortexing for 5 s and centrifuging for 1 min at 500 rpm. The recovered DNA in water, denoted as “unsort”, was saved for later measurement of enrichment factor.

### Dielectrophoretic sorting

The thermocycled drops were transferred to a 1 ml syringe and reinjected to a microfluidic dielectrophoretic (DEP) sorter (Fig. 2) at 50 μl/h^8,20^. The syringe was placed vertically so that the drops remained at the top and closely packed. Individual drops were separated after entering the sorter by a spacer oil of HFE-7500 with a flow rate of 950 μl/h. Another stream of HFE-7500 oil at 1000 μl/h was introduced at the sorting junction to drive the drops to waste collection when the DEP force was off. A syringe at -1000 μl/h was used to produce a negative pressure at the waste collection to further ensure unsorted drops flowed to waste. The salt water electrodes and moat shielding were filled with 2M NaCl solution. A laser of 100 mW, 532 nm was focused upstream of the sorting junction to excite droplet fluorescence. Photomultiplier tubes (PMTs, Thorlabs, PMM01 model) were focused on the same spot to measure emission fluorescence. A data acquisition card (FPGA card) and a LabVIEW program (available at GitHub: https://github.com/AbateLab/sorter-code) (National Instruments) were used to collect PMT outputs and activate the salt electrode when the emission fluorescence intensity is higher than a pre-set threshold. A high-voltage amplifier (Trek) was used to amplify the electrode pulse to 0.8-1 kV for DEP sorting. The sorted drops were collected into a 1.5 ml Eppendorf DNA LoBind tube.

### DNA recovery and 2^nd^ round of enrichment

DNA from sorted drops was recovered by breaking the emulsion with 10% (v/v) solution of perfluoro-octanol (Sigma-Aldrich, catalog no. 370533) and addition of 20 μl DI water, followed by gentle vortexing for 5 s and centrifugation for 1 min at 500 rpm. 2 μl of the recovered DNA, denoted as “single sort”, was saved for later measurement of the enrichment factor by qPCR. The remaining 18 μl recovered DNA was processed with a 2^nd^ round of droplet TaqMan PCR and DEP sorting as described above. After sorting, the sorted drops were broken and the recovered DNA, denoted as “double sort” used to measure the degree of enrichment.

### Quantitative PCR analysis of sorted droplets

We used a multiplex TaqMan PCR, with one FAM based probe targeting ΦX 174 DNA and one Cy5 based probe targeting lambda DNA to quantify ΦX 174 and lambda DNA in “unsort”, “single sort” and “double sort”. The PCR reaction was set as: 1X Platinum Multiplex PCR Master Mix, 200 nM TaqMan probes, 1 μM forward primers and 1 μM reverse primers (IDT), recovered DNA and DNAse-free water to bring the volume to 25 μl. The PCR was performed in a QuantStudio 5 Real-Time PCR System (Thermo Fisher Scientific) using the following parameters: 95°C for 2 min; 40 cycles of 95°C for 30 s, 60°C for 90 s and 72°C for 30 s. C_t_ values for each sample were obtained and used to compute the enrichment factor. All primer and TaqMan probe sequences are listed in Supplementary Table S1. The TaqMan assays were tested for specificity and linearity by constructing a serial dilution of ΦX 174 DNA with a fixed concentration of lambda DNA. We obtained two C_t_ values for ΦX 174 (FAM) and lambda (Cy5) for each of the “unsort”, “single sort” and “double sort” samples, to compute the enrichment factors for each round of sorting.

### HIV associated DNA sample preparation

The HIV infected cells were prepared by plating resting CD4 T cells from an ART treated person at ∼1 infected cell per 5 wells (∼100 total cells per well), followed by stimulation and a period of *in vitro* culture to allow proliferation^28^. Non-HIV infected Jurkat cells (ATCC® TIB-152™) were cultured following the provided protocol. DNA were extracted from clonally expanded cells (from one well of the culture plate) and from Jurkat cells using Quick-DNA™ Miniprep Plus Kits (Zymo research, catalog no. D4068) according to the manufacturer’s instructions, and mixed at a 1:30 ratio.

### Compound enrichment of single HIV genomes

The DNA mixture was processed with droplet TaqMan PCR and 1^st^ DEP sorting as described above using HIV *pol* specific TaqMan probe and biotinylated primers. All primer and TaqMan probe sequences for HIV are listed in Supplementary Table S2. The sorted emulsions were broken using perfluoro-1-octanol and the aqueous was diluted in 5 μl H_2_O. The aqueous layer containing sorted DNA was then added to streptavidin conjugated magnetic beads (Dynabeads MyOne Streptavidin C, Thermo Fisher Scientific) and incubated for 15 min. D1 buffer from REPLI-g single cell kit (Qiagen, catalog no. 150343) was added to denature the DNA. Biotinylated primers and amplicons were attached to the magnetic beads and removed after transferring supernatant to a fresh tube. The MDA reaction mixture was then prepared with a REPLI-g single cell kit by following the manufacturer’s protocol and emulsified by a flow-focus droplet maker (diameter ∼20 μm) as described above. The emulsion was collected to a 1 ml syringe and incubated at 30 °C for 20 h. After incubation, MDA droplets and 2^nd^ TaqMan PCR reagents were injected into a microfluidic merger device^26^. PCR reagent drops were formed on chip and merged with MDA drops pairwise. Merging was achieved at a salt electrode connected to a cold cathode fluorescent inverter and DC power supply (Mastech) to generate a ∼2 kV AC signal from a 2 V input voltage. The merged drops (diameter: 40 μm) were collected to PCR tubes. The bottom oil layer was removed and replaced with FC-40 fluorinated oil with 5% (w/w) PEG-PFPE surfactant for thermal cycling: 3 min at 86 °C; 35 cycles of 30 s at 86 °C, 90 s at 60 °C and 30 s at 72 °C; and finally, 5 min at 72 °C. After PCR, the drops were reinjected into a DEP sorter for the 2^nd^ sorting round as described above.

### Library preparation and sequencing from sorted droplets

The sorted single droplets with their carrier oil were collected into individual PCR tubes and dried out in a vacuum chamber. 1 μl DI H_2_O was added to dissolve the sorted DNA. The dissolved DNA was then tagmented using 0.6 μl TD Tagmentation buffer and 0.3 μl ATM Tagmentation enzyme from Nextera DNA Library Prep Kit (Illumina, catalog no. FC-121-1030) for 5min at 55 °C. 1 μl NT buffer were added to neutralize the tagmentation. The tagmented DNA was then mixed with PCR solution containing 1.5 μl NPM PCR master mix, 0.5 μl of each index primers i5 and i7 from Nextera Index Kit (Illumina, catalog no. FC-121-1011) and 1.5 μl H_2_O, and placed on a thermal cycler with the following program: 3 min at 72 °C; 30 s at 95 °C; 20 cycles of 10 s at 95 °C, 30 s at 55 °C, and 30 s at 72 °C; and finally 5 min at 72 °C. The DNA library was purified using a DNA Clean & Concentrator-5 kit (Zymo Research, catalog no. D4004), size-selected for 200-600 bp fragments using Agencourt AMPure XP beads (Beckman Coulter), and quantified using Qubit dsDNA HS Assay Kit (Thermo Fisher Scientific) and High Sensitivity DNA Bioanalyzer chip (Agilent). The library was sequenced using Illumina Miseq and ∼1 million paired-end reads of 150 bp were used for each sorted droplet. Sequencing reads were mapped to the HIV reference genome (HXB2) using Bowtie 2^33^. Genomic coverage as a function of genome position was generated using SAMtools^34^. The non-HIV regions of chimeric reads were extracted using extractSoftclipped (https://github.com/dpryan79/SE-MEI) and analyzed by a web base tool for integration sites (https://indra.mullins.microbiol.washington.edu/integrationsites/).

## Acknowledgements

We thank James I. Mullins at University of Washington for providing HIV infected cells. We also thank members of the Abate lab, in particular Leqian Liu, Cyrus Modavi, David J. Sukovich and Samuel C. Kim for helpful discussions. This work was supported by the Chan Zuckerberg Biohub, the National Institutes of Health (NIH) (Grant No. R01-EB019453-01, R01-HG008978-01 and DP2-AR068129-01), the National Science Foundation CAREER Award DBI-1253293.

