## Supplemental data for "Sequencing ultra-rare targets with compound nucleic acid cytometry"

### Derivation of equations for enrichment power

We define the enrichment power as the ratio of target purity in the sample after enrichment to before,

$$\text{Enrichment power} = \frac{\text{Target purity after enrichment}}{\text{Target purity before enrichment}},$$

where $N_T$ is the number of target molecules, $N_O$ is the total number of off-target molecules, $D$ is the total number of droplets and $f$ is the false positive rate of the assay. In the mixed DNA sample, the total number of molecules is $N_T + N_O$ and thus

$$\text{Target purity before enrichment} = \frac{N_T}{N_T+N_O},$$

We encapsulate these $N_T + N_O$ molecules into $D$ droplets, such that each drop contains $\frac{N_T+N_O}{D}$ molecules. The DNA mixture is partitioned at limiting dilution such that individual droplets rarely contain more than one target expected by Poisson statistics, thus $N_T$ drops are PCR positive. We also expect $fD$ false positive drops. Because the false positive rate is generally small and the target is ultra-rare, the instances of false positive and true positive in the same droplet are infrequent. In together, $N_T + fD$ drops are sorted and the number of molecules in the sorted sample is $\frac{N_T+N_O}{D}(N_T + fD)$, of which $N_T$ molecules are the targeted ones. Therefore,

$$\text{Target purity after enrichment} = \frac{N_T}{\frac{N_T+N_O}{D}(N_T+fD)},$$

and to simplify the enrichment power to

$$\text{Enrichment power} = \frac{\frac{N_T}{\frac{N_T+N_O}{D}(N_T+fD)}}{\frac{N_T}{N_T+N_O}} = \frac{1}{\frac{N_T}{D}+f}$$

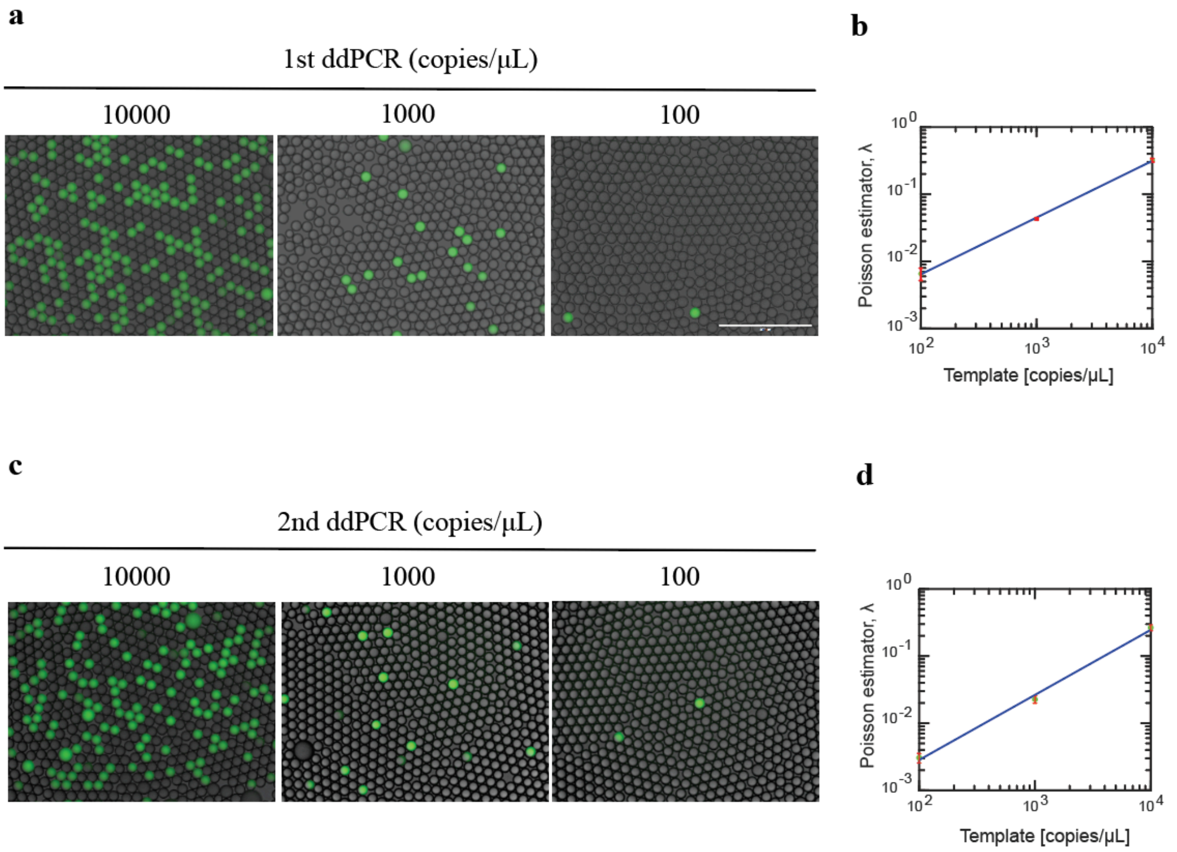


Supplementary Fig. 1 TaqMan sets reliably detect ΦX 174 DNA in compound NAC. Fluorescence images of droplets after PCR amplification with TaqMan sets used in **(a)** $1^{st}$ NAC and **(c)** $2^{nd}$ NAC for ΦX 174 DNA templates at varying concentrations. The template copy number per droplet estimated by assuming a Poisson distribution scales with the input template concentrations during ddPCR in **(b)** $1^{st}$ NAC, and **(d)** $2^{nd}$ NAC.

Supplemental Table S1: Primers and probes for compound enrichment of ΦX 174 virus DNA

| Primer Name | Sequence (5→3') |
|---|---|
| 1st NAC | |
| ΦX 174 probe1 | /56-FAM/TAAATCGAA/ZEN/GTGGACTGCTGGCGG/3IABkFQ/ |
| ΦX 174 F1 | GCAGGAATTACTACTGCTTGTTTAC |
| ΦX 174 R1 | GAATCGTTAGTTGATGGCGAAAG |
| 2nd NAC | |
| ΦX 174 probe2 | /56-FAM/CGTATGCAG/ZEN/GGCGTTGAGTTC/3IABkFQ/ |
| ΦX 174 F2 | GCAGATGGATAACCGCATCA |
| ΦX 174 R2 | CCTTATGGCCGTCAACATACA |
| Post sorting qPCR | |
| ΦX 174 probe3 | /56-FAM/ATGGAACTG/ZEN/ACCAAACGT/3IABkFQ/ |
| ΦX 174 F3 | GCGCTCTAATCTCTGGGCAT |
| ΦX 174 R3 | CAAAGAAACGCGGCACAGAA |
| lambda probe3 | /5Cy5/TGAGGTGCT/TAO/TTATGACTCTGCCGC/3IAbRQSp/ |
| lambda F | GCGAGTATCCGTACCATTCAG |
| *lambda* R | TCCCTTTCGGCATACCATTT |

Supplemental Table S2: Primers and probes for compound enrichment of HIV provirus DNA

| Primer Name | Fluorophore, quencher | Sequence (5→3') |
|---|---|---|
| 1st NAC | | |
| *pol* probe | FAM, ZEN-3'IBFQ | AAGCCAGGAATGGATGGCC |
| *pol* F | | /5Biosg/ GCACTTTAAATTTTCCCATTAGYCCTA |
| *pol* R | | /5Biosg/ CAAATTTCTACTAATGCTTTTATTTTTTC |
| 2nd NAC | | |
| LTR probe | FAM, ZEN-3'IBFQ | CTGGTAACTAGAGATCCCT |
| LTR F | | GCCTCAATAAAGCTTGCCTTGA |
| LTR R | | GCTAGAGATTTTCCACACTGACTARA |
| *tat probe* | Cy5, TAO-3'IBRQSp | CTATGGCAGGAAGAAGCGGAGACAGC |
| *tat* F | | TGTAAAAAGTGTTGCYTTCATTG |
| *tat* R | | ACTACTTACTGCTTTGRTAGAG |
